# Swung and spun in weightlessness : Evidence of immediate canalar underdetection of rotations in parabolic flight

**DOI:** 10.64898/2026.06.30.735470

**Authors:** Tess Bonnard, Emilie Doat, Dominique Guehl, Etienne Guillaud

## Abstract

Despite extensive research on vestibular function in microgravity, particularly during orbital and parabolic flight exposure, several gaps remain regarding the spontaneous behavior of vestibular organs under non-terrestrial gravitoinertial conditions. In particular, semicircular canal function, typically assessed through vestibulo-ocular reflex (VOR) recordings, has yielded inconsistent findings, with reports describing either no effect or reduced performance in microgravity. Moreover, many of these studies are limited by methodological constraints that reduce the interpretability of their conclusions.

To clarify these discrepancies, we evaluated horizontal and vertical VOR responses during parabolic flights to assess semicircular canal function under transient weightlessness. Participants were passively rotated at a constant frequency and amplitude during normogravity and microgravity phases, centered along the head’s vertical or inter-aural axis. Eye movements were recorded binocularly using infrared eye-tracking in darkness to eliminate visual influences, while participants were tightly restrained to minimize proprioceptive variability.

Results show a reduction in VOR gain during microgravity in both axes, despite consistent rotational stimulation across gravity conditions. In addition, VOR gain remained reduced after parabolas in the horizontal plane, whereas vertical VOR performance was preserved.

These are the first results to demonstrate an immediate alteration of semicircular canal function in weightlessness. Possible sources of the reduction in VOR performance in 0g are discussed. We also propose that the observed post-flight effects reflect a down-weighting of semicircular canal inputs during multisensory integration.

## INTRODUCTION

Gravity is omnipresent on Earth and is continuously detected by the vestibular system. This sensory system comprises the otolith organs, which enable the detection of the constant terrestrial gravitoinertial vector (approximately 1 g). Recent studies have shown that the otolithic system remains stable in detecting translational acceleration even when gravitational forces are absent (Bonnard et al., 2026a). In addition, the vestibular system includes three semicircular canals in each ear, which function as biological gyroscopes to detect angular rotations of the head. However, the ability of the canalar system to accurately detect rotation in weightlessness remains unclear, despite extensive experimental studies yielding controversial results. Of particular importance is how the vestibular system maintains coherence during the initial exposure to weightlessness, as this system is thought to be a primary source of space motion sickness – a condition affecting approximately 70% of astronauts (Davis et al., 1988b).

The vestibulo-ocular reflex (VOR) is commonly used to assess the integrity and function of the semicircular canals. This reflex relies on a three-neuron arc and stabilizes gaze during head movements by generating compensatory eye movements in the direction opposite to head motion. Anatomical and functional studies in animal models have shown that the angular VOR is predominantly driven by semicircular canal inputs, with only minimal contributions from the otolith organs (Fetter, 2007).

Previous studies conducted during orbital flight have reported that horizontal VOR gain remains unchanged following long-duration exposure to weightlessness (Berthoz et al., 1986; Benson & Vieville, 1986; Thornton et al., 1989; Grigorova & Kornilova, 1996; Clarke et al., 2000). However, other investigations have yielded contradictory findings, including reductions in the horizontal gain of the oculomotor compensation for active head rotation in the early days of flight for three astronauts (Clément et al., 2019) or in the hours following return to Earth (Clément et al., 2019; Clarke et al., 2000). Only a limited number of studies have examined VOR responses in parabolic flight (PF). Oman et al. (1996) found no change in the gain of vestibular-induced nystagmus in this analog, while Lackner and Graybiel (1981) observed a reduction in the intensity of post-rotatory nystagmus under microgravity.

Several methodological considerations must be taken into account when interpreting these findings. In orbital flight studies, VOR assessments mostly rely on dynamic, voluntary head movements performed by participants rather than passive rotations (Berthoz et al., 1986; Benson & Vieville, 1986; Thornton et al., 1989; Grigorova & Kornilova, 1996; Clarke et al., 2000; Clément et al., 2019). Such approaches introduce proprioceptive inputs and cognitive influences (e.g. efference copy), which may confound the interpretation of vestibular reflex responses (Doerr et al., 1981; Jurgens & Mergner, 1989). During voluntary head movements, cervico-ocular reflexes – initiated by neck proprioception – enable accurate eye movement on their own (Bronstein and Hood, 1986). Moreover, the effects of changes in gravitoinertial load on these cervico-ocular reflexes remain poorly understood. In addition, in orbital flight studies, measurements are typically obtained only after four or more days of weightlessness, by which time space motion sickness has subsided and adaptive processes are likely to have occurred. As a result, most reported outcomes may reflect adaptive changes rather than the immediate functional state of the reflex during initial exposure to microgravity.

In contrast, parabolic flight experiments can be conducted before adaptive processes occur. However, such studies remain limited in number, and their methodologies are also open to criticism. The study by Oman et al. (1996) relied on passive rotations of participants who were administered scopolamine–dexedrine, a combination known to affect vestibular function (Weerts et al., 2015; Bestaven et al., 2016), potentially confounding the interpretation of VOR measurements. Furthermore, in two studies (Oman et al., 1996; Lackner & Graybiel, 1981), the protocol involved measuring nystagmus following the abrupt cessation of whole-body rotation. The results were derived from assessments of velocity storage—a perceptual process modulated by additional vestibular stimulation (the “dumping effect”; Oman & Balkwill, 1993). In these experiments, participants underwent rotations during parabolic flight, which subjected them to gravitational acceleration transitions from 1g to 0g, including a 1.8g phase. As a result, the degree to which these findings reflect human vestibular responses to weightlessness remains unclear.

Understanding whether the semicircular canals continue to function normally during the initial phase of microgravity exposure is critical in the context of the sensory conflict theory of motion sickness. As originally proposed by Reason (1978), motion sickness arises when sensory signals from different sources—either between systems (inter-sensory conflict) or within a single system (intra-sensory conflict)—are incongruent, thereby complicating multisensory integration and perceptual interpretation. In the case of space motion sickness (SMS), an otolitho-canalar conflict has been hypothesized as a primary underlying mechanism (Reason, 1978; Oman, 1998). A potential source of conflict could arise if vestibular capacities were influenced by the gravitational context, leading to inconsistencies with other proprioceptive or visual sensory cues. Furthermore, if one vestibular subsystem— whether otolithic or canal-based—were specifically altered by changes in gravity while the other remained unaffected, this imbalance could generate a nauseogenic otolitho-canalar conflict during any head movement. Based on previous experiments (Bonnard et al., 2026), we suggested that otolithic function is unaffected by either weightlessness or hypergravity. Therefore, it is critical to assess whether semicircular canal function is preserved or disrupted during the initial transition to microgravity. Such insights are essential for advancing our understanding of the mechanisms that underlie the onset of space motion sickness (SMS).

To determine whether semi-circular canals function is preserved or modulated by changes in the gravitoinertial environment, we assessed the VOR during parabolic flight using passive full body rotations in both yaw and pitch, in darkness. This approach was designed to provide controlled vestibular stimulation under alternating normogravity and microgravity conditions, thereby assessing the effect of gravitational load on VOR responses while minimizing the influence of adaptation or fatigue. We hypothesized that VOR performance in weightlessness would be altered in the vertical plane, where otolithic inputs are involved, whereas horizontal VOR responses would remain either unchanged, as suggested by previous literature. In addition, baseline measurements obtained in normogravity before the experimental protocol were compared with post-parabolas responses. This approach, combined with parallel monitoring of participant discomfort, allowed us to investigate potential relationships between space motion sickness susceptibility, symptom severity, and changes in semi-circular canals function.

## MATERIAL & METHODS

### Participants

Twelve participants were enrolled in this protocol (mean age: 25 ± 5 years; 5 females, 6 males). Before the flight, all participants completed a medical questionnaire, which was reviewed and approved by an aerospace physician. This evaluation included an effort electrocardiogram performed and analyzed by a cardiologist. No history of sensory disorders was reported. Prior to participation in the present study, all subjects provided written informed consent following an inclusion visit and a mandatory cooling-off period. Ethical approval for this study was granted by the national Ethics Committee «Comité de protection des personnes Ile de France II» n°2023-A02145-40 and all experiments were conducted according to Helsinki Agreement.

### Protocol

Eleven out of twelve participants performed two passive vestibulo-ocular reflex (VOR) tests. One participant was unable to perform the test during the parabolic flight session because of severe motion sickness. Two original experimental setups were developed to generate manually induced passive rotations: one around a vertical axis (yaw rotations) and one around a horizontal axis (pitch rotations). For clarity, these setups are hereafter referred to as the “rotating-chair” (yaw rotation) and the “swing-chair” (pitch rotation). Both setup were attached to the aircraft floor.

The rotating-chair consisted of a seat mounted on a single ball bearing aligned with the midpoint between the participant’s ears (Figure 1, right). A race car–type seat equipped with a seatbelt was used to ensure comfort and safety, while maintaining trunk stabilization across all gravity conditions. A handle positioned at shoulder width and mid-arm height of experimenter behind the seat enabled manual oscillation of the system.

**Figure 1.**
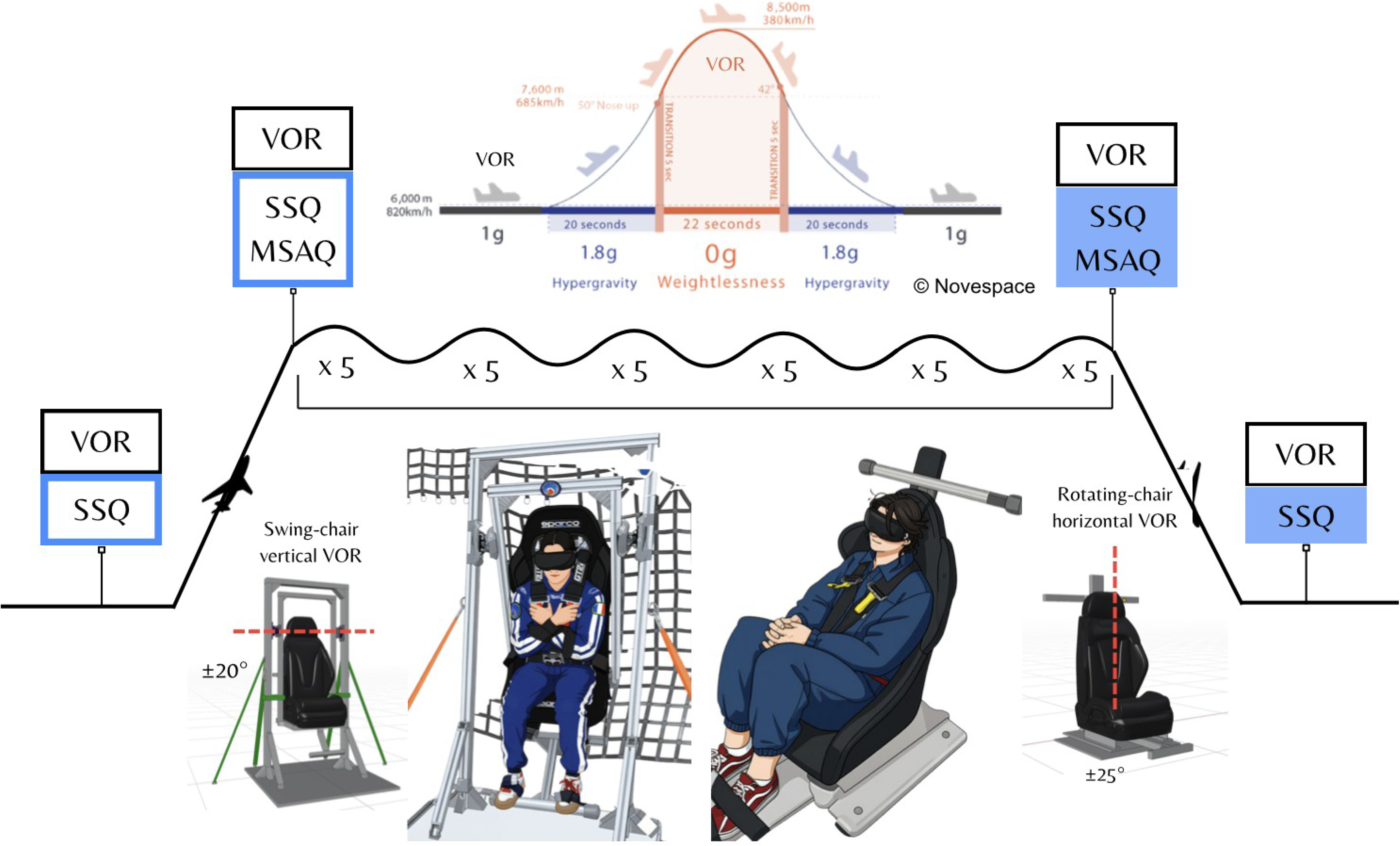
Overview of the experimental protocol. The top panel illustrates the sequence of parabolic maneuvers. The middle panel shows the flight profile and the timing of measurement sessions. The bottom panel presents the two experimental setups. Light blue frames indicate pre-flight measurements (Pre-Ground and Pre-Flight), whereas light blue filled markers indicate post-flight measurements (Post-Flight and Post-Ground). Schematic representations of the experimental setups illustrate the two setups during parabolic flight under real operating conditions and are representative of participant positioning. Schematic renderings of the setups, generated using Blender, are also shown to illustrate the experimental structures and rotation axes (dashed red lines). SSQ: Simulator Sickness Questionnaire, MSAQ: Motion Sickness Assessment Questionnaire.

The swing-chair consisted of a race car seat mounted on a rigid swing frame, attached to an arch with a pitch-axis rotation pivot located at the participant’s ear level (approximately 130 cm from the floor; Figure 1, left). Handles positioned at multiple vertical heights along the arch and behind the seat enabled manual control of pitch-axis rotations.

VOR testing was performed using the Pupil Labs Neon eye-tracking system (Pupil Labs, Neon “The googles, they do everything”, Berlin, Germany). This device is equipped with two infrared cameras that record pupil movements of each eye, as well as inertial measurement units (IMUs) to capture head motion. The visor of the headset was blackened to eliminate visual input and allow VOR assessment in complete darkness. Participants were tightly strapped to the seat to preserve proprioceptive clues in weightlessness. All recordings were initiated and stored using the Neon Companion application. Interpupillary distance was measured using a calibrated scale and entered into each participant’s profile within the recording application prior to testing. Before each trial, real-time visualization of eye positions from the infrared cameras was checked to ensure proper alignment and centering of both eyes within the field of view.

Participants were instructed to initially close their eyes, center their gaze straight ahead, and then reopen their eyes while maintaining this neutral position, without vergence. They were instructed to use their initial eye position as a spatial reference and to compensate for seat motion to maintain their gaze in this allocentric direction. Although the task was challenging, participants were encouraged to minimize conscious effort and allow their eyes to respond naturally.

For the rotating-chair setup, rotational amplitude was fixed at 50° (±25° from the starting position), with oscillations at 0.5 Hz. For the swing-chair setup, rotations were set to 40° (±20° from the starting position), at 0.54 Hz, matching the system’s natural oscillation frequency at 1 g. These parameters were consistent with previous literature and were selected for their reproducibility during pre-testing under normogravity conditions.

Parabolic flights were conducted during the VP190 campaign organized by the Centre National d’Études Spatiales (CNES), aboard the Airbus A310 Zero-G operated by Novespace in Mérignac (France). Each flight comprised 31 parabolic maneuvers, following a sequence of altered gravity phases (Figure 1) : hypergravity (1.8 g, 24s), microgravity (0 g, 22s), and a return to hypergravity (1.8 g, 20s), interspersed with periods of normogravity (1g) during steady flight. Each parabola lasted approximately 1 minute, with 2-minute intervals between successive parabolas. The total flight duration ranged from 2.5 to 3 hours, including takeoff, transit to the flight zone, the parabolic session, return, and landing. To prevent motion sickness (MS), scopolamine is commonly administered prior to flight, as it is known to attenuate vestibular processing (Weerts et al., 2015; Bestaven et al., 2016). However, in the present study, no medication was administered in order to preserve vestibular function and maintain participant alertness.

Baseline recordings of 1-minute duration were conducted in normogravity at four distinct time points: before takeoff while on the airport tarmac (Pre-Ground), after takeoff but immediatly prior to the initiation of the parabolic session (Pre-Flight), immediately following the last parabola (Post-Flight), and after landing (Post-Ground). Experimental trials included 0g trials, which were performed during the 20s periods of weightlessness within each parabola, as well as comparative normogravity trials conducted between parabolas. This approach helped avoid longitudinal biases (such as fatigue, sickness, or reduced concentration) that could affect the comparison of gravitational effects. Throughout the hypergravity phases and transitions, participants were motionless and instructed to remain securely fastened to ensure data consistency and safety.

Each participant completed a total of 10 trials for both the vertical and horizontal VOR tasks. The trials were organized into sessions of 5 trials each, with the order of tasks (vertical or horizontal) alternated and counterbalanced across participants. Specifically, half of the participants began with 5 horizontal VOR trials, while the other half started with 5 vertical VOR trials. During the flight, each participant performed one session of 5 trials for each VOR orientation (vertical and horizontal) in the first half of the flight, and another session of 5 trials for each orientation in the second half. In total, participants were engaged in the task for 20 parabolas and had 11 parabolas designated as rest periods. On average, and excluding baseline measurements, each participant performed a large number of rotations in each task (horizontal VOR: 197 ± 103 rotations at 1g, 159 ± 64 at 0g; vertical VOR: 218 ± 79 rotations at 1g, 177 ± 67 at 0g).

### Data analysis

Recorded data were stored on smartphones (one per setup) and transferred to Pupil Cloud (Pupil Labs, Berlin, Germany) after completion of the experimental protocol. This transfer enabled a posteriori visualization of eye recordings and extraction of raw data at 200 Hz, corresponding to the acquisition frequency of both the infrared cameras and inertial measurement units (IMUs). Vertical and horizontal eye positions (in degrees), as well as head yaw and pitch velocities were extracted from Pupil Cloud.

All subsequent analyses were performed using MATLAB R2024b (MathWorks, Natick, MA, USA). Head motion velocity signals were filtered using a zero-phase fourth-order Butterworth band-pass filter (0.1–3 Hz), corresponding to the typical frequency range of physiological head movements. Angular positions were subsequently obtained by integrating the filtered signals. Head rotations were identified when peak head velocity exceeded 40°·s⁻¹, a threshold empirically selected to distinguish passive head movements from noise. The onset and offset of each head rotation were determined by the first velocity zero-crossings before and after the peak velocity, respectively. Eye movement data were preprocessed to remove blinks and saccades, defined as events exceeding 100°·s⁻¹, to reduce noise and isolate the slow-phase eye movements associated with the VOR.

Amplitude, duration, angular velocity peak, and median were computed for each head motion. VOR gains and latencies were calculated separately for each head displacement on the left, on the right, up or down. Two complementary approaches were used: (i) amplitude gain, defined as the ratio of eye displacement amplitude to head displacement amplitude, and (ii) velocity gain, defined as the ratio of median eye velocity to median head velocity, both calculated during the head movement period. Latency was defined as the temporal lag between eye and head position signals, estimated using a cross-correlation method. Median eyes positions of each movement were analyzed in both the horizontal and vertical planes for each experimental setup to assess potential deviations induced by changes in gravity level. To simplify further analysis, and given the verified symmetrical behavior between the left and right eyes, data from both eyes were averaged for the statistical analysis.

Motion sickness was assessed using standardized scales and questionnaires. The Simulator Sickness Questionnaire (SSQ; Kennedy et al., 1993) was administered to each participant to evaluate their current symptoms at four time points: Pre-Ground, Pre-Flight, Post-Flight, and Post-Ground. The SSQ comprises 16 items assessing both nausea-related symptoms (e.g., stomach awareness, burping, sweating) and oculomotor disturbances (e.g., eye strain, headache, blurred vision). The Motion Sickness Assessment Questionnaire (MSAQ ; Gianaros et al., 2001) was additionally administered during the Pre-Flight and Post-Flight sessions, as it captures symptom dimensions complementary to those assessed by the SSQ. The MSAQ includes 16 items rated on a 9-point scale (from 1, “not at all,” to 9, “severely”). It provides subscale scores across four domains: gastrointestinal, sopite-related, peripheral, and central symptoms.

Additionally, motion sickness was monitored after each parabola throughout the entire parabolic session using the Motion Sickness Severity Scale (Czeisler et al., 2023). This scale comprises 7 levels from no symptoms (0) to vomiting (6) to enable continuous tracking of subjective symptom progression during flight. To enable correlations with facial skin color measurements, acquired during the breaks, MSSS ratings were calculated as the average of the two preceding and two subsequent parabolas surrounding each break (i.e., every five parabolas) Facial skin color was measured using a colorimeter (ColorReader EZ, DC10-3, Datacolor GmbH, Marl, Germany). Placed on the cheek at a fixed position, this device captures color directly on the skin surface and illuminating it with an integrated LED, thereby avoiding interference from ambient light. The RGB color scale was used for analysis and normalized individually for each participant: each color component (red, green, blue) was expressed as a percentage of the baseline measured before the experiment.

### Statistics

Repeated-measures analyses of variance (ANOVA) with Tukey (HSD) post hoc were conducted to assess the effects of Gravity, Direction, the Gravity × Direction interaction, Pre/Post condition, Flight vs. Ground condition, and the Pre/Post × Flight/Ground interaction for both experimental setups. Prior to conducting the ANOVA, homogeneity of variance was assessed using Levene’s test and was confirmed for all analyses (p > 0.05). Sphericity was evaluated using Mauchly’s test of sphericity. When the sphericity assumption was violated (p < 0.05), corrections were applied according to the epsilon (ε) value: the Greenhouse–Geisser correction was used for ε < 0.75, whereas the Huynh–Feldt correction was applied for ε ≥ 0.75. Effect sizes were reported as partial eta squared (η²), calculated as the ratio of the effect sum of squares to the sum of the effect and error sum of squares. Results are expressed as mean ± standard error (SE).

## RESULTS

### Gravity Effects on the VOR

For the *horizontal test*, head movement parameters (amplitude, velocity, and duration) did not vary as a function of gravity level (Amplitude: 0 g = 52.2 ± 1.4°, 1 g = 52.9 ± 0.9°, F(1,10) = 0.43, p = 0.53; Velocity Peak: 0 g = 78.5 ± 2.2°·s⁻¹, 1 g = 79.1 ± 1.5°·s⁻¹, F(1,10) = 0.13, p = 0.73; Duration: 0 g = 0.99 ± 0.003s, 1 g = 0.99 ± 0.003s, F(1,10) = 0.14, p = 0.72). This confirms that experimenter-generated passive rotations were reproducible and comparable between normogravity and microgravity conditions (Figure 3B-C). In the same way, stimulations were symmetrical, with no significant differences between left and right (Amplitude: Right = 52.5 ± 1.17°, Left = 52.6 ± 1.17°, F(1,10) = 0.22, p = 0.65; Velocity Peak: Right = 79.3 ± 1.88°·s⁻¹, Left = 78.3 ± 1.87°·s⁻¹, F(1,10) = 1.00, p = 0.34; Duration: Right = 0.98 ± 0.003 s, Left = 0.99 ± 0.003 s, F(1,10) = 3.67, p = 0.084).

**Figure 2.**
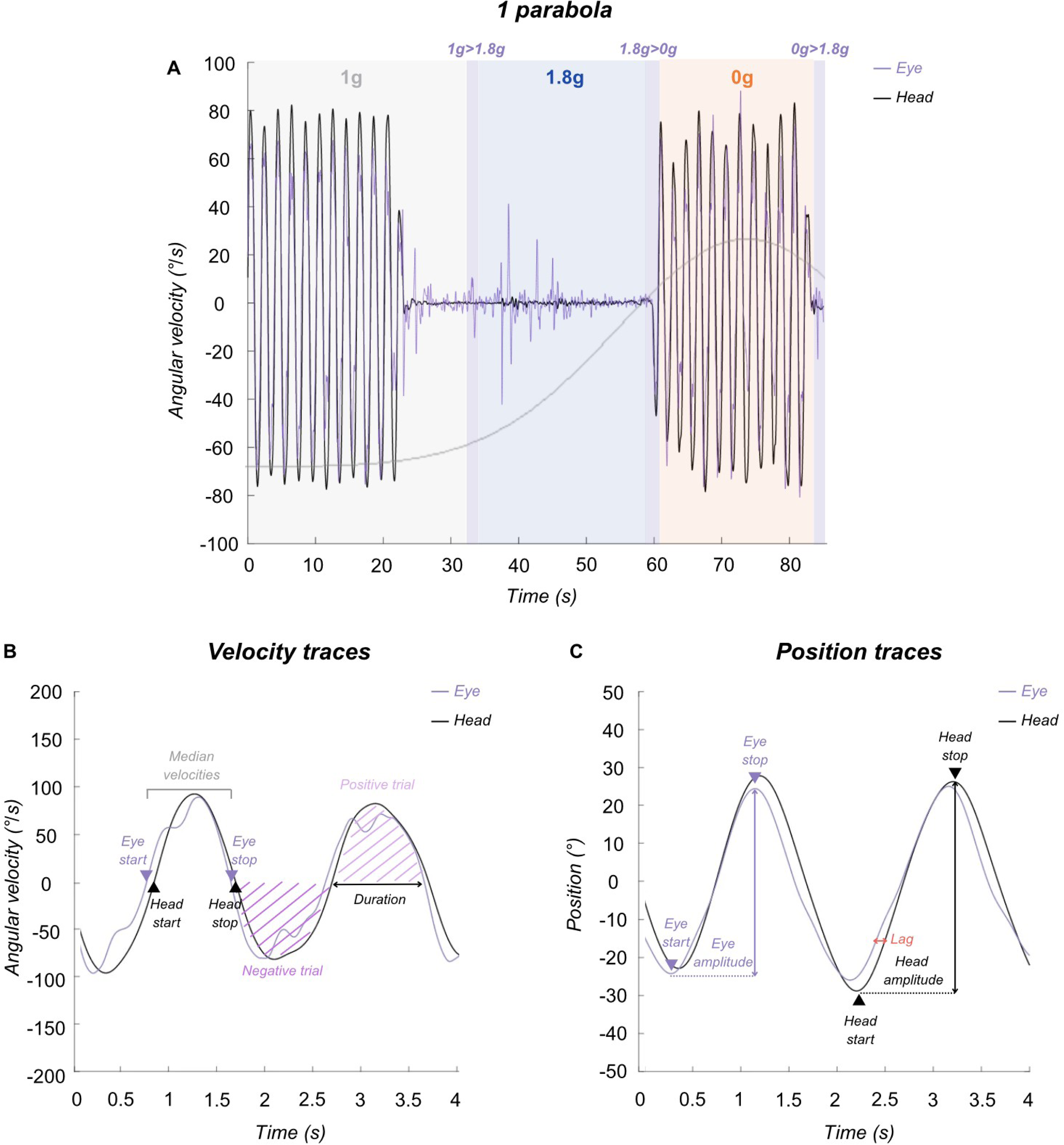
Data overview. Representative traces from a single participant recorded using the rotating-chair. Eye movements are depicted in purple and head movements in black. Panel A illustrates an example of a single trial with eye and head velocities, including phases of normogravity (grey), microgravity (orange), and the hypergravity transition period (blue), interspaced with transition periods (purple). Participant was rotated during the 1g period before plane’s nose-up maneuver and during the microgravity period. Panels B and C present the velocity and position traces of two head movements. In both panels, the onset and offset of head and eye movements are indicated.

**Figure 3.**
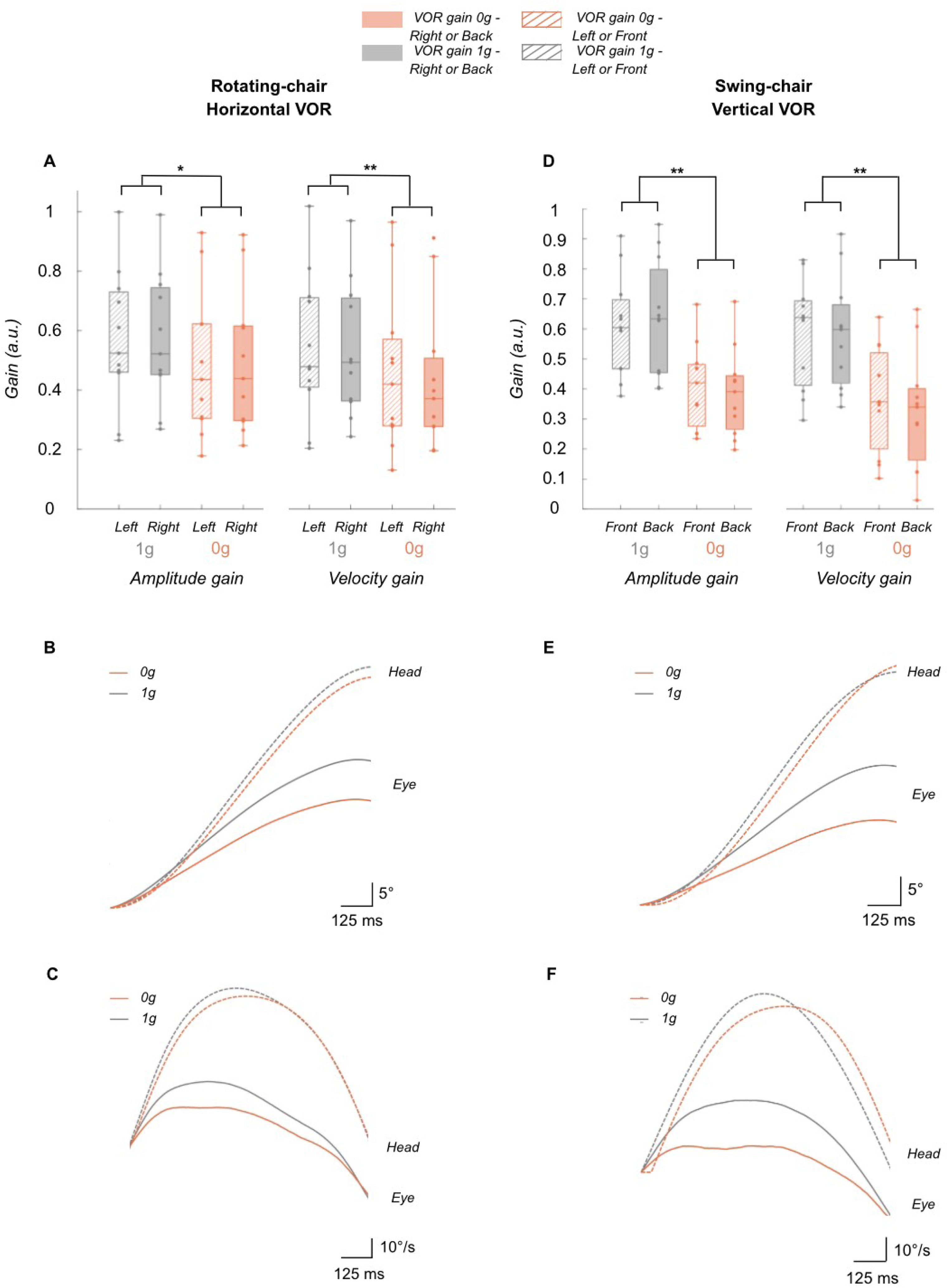
Horizontal (left panels, A-B-C) and vertical (right panels, D-E-F) VOR gains depending on gravity. In boxplots (A & D), microgravity is represented in orange and normogravity in flight in grey. Dashed orange represents backward or leftward motion whereas filled orange depicts forward or rightward motion, depending in the experimental setup used. Grand averages across participants of eye and head positions (B & E) and velocites (C & F) are presented in each gravity condition. Panels B and E represent eye and head positions (°) whereas panels C & F represent eye and head velocites (°/s) Repeated-measures ANOVA with Tukey post-hoc. *, p<0.05, ** p<0.001.

Eye movement amplitudes were significantly smaller (17%) in microgravity compared to normogravity (1 g: 30.43 ± 2.73°; 0 g: 25.14 ± 2.56°; F(1,10) = 17.33, p = 0.002, η²ₚ = 0.63) (Figure 3B). This resulted in significantly lower amplitude gain in microgravity (1 g: 0.57 ± 0.05; 0 g: 0.49 ± 0.05; F(1,10) = 8.31, p = 0.016, η²ₚ = 0.45; Figure 3A). There was also a non-significant tendency toward lower eye peak velocities in microgravity (1 g: 64.58 ± 3.98°·s⁻¹; 0 g: 59.59 ± 3.80°·s⁻¹; F(1,10) = 4.11, p = 0.07, η²ₚ = 0.29) (Figure 3C), leading to a significant decrease in velocity gain in microgravity (1 g: 0.54 ± 0.05; 0 g: 0.45 ± 0.05; F(1,10) = 12.90, p = 0.005, η²ₚ = 0.56) (Figure 3A). Eye peak velocity also showed a significant directional effect, with faster movements during rightward rotations (rightward: 64.13 ± 3.92°·s⁻¹; leftward: 60.03 ± 3.89°·s⁻¹; F(1,10) = 12.67, p = 0.005, η²ₚ = 0.56).

For the horizontal VOR, the eye position phase lead relative to head motion was not affected by gravity (1 g: -7.02 ± 5.74 ms; 0 g: -11.48 ± 7.07 ms; F(1,10) = 2.52, p = 0.15). Additionally, the median vertical eye position remained stable regardless of gravity conditions (1 g: 0.27° ± 0.04°; 0 g: 0.24 ± 0.05°; F(1,10) = 0.13, p = 0.73).

Regarding the *vertical VOR*, slight but significant changes in head kinematics were observed between the two gravitational conditions and between the forward and backward directions. In microgravity, where kinematics were strictly driven by the experimenter’s hand actions, head displacement amplitude was significantly 6% larger (1 g = 39.41 ± 0.44°, 0 g = 41.70 ± 0.41°; F(1,10) = 53.85, p < 0.001, η²ₚ = 0.84) (Figure 3E). This effect was accompanied by a directional effect, with backward sway being 0.7% larger than forward sway (backward = 40.37° ± 0.50°, forward = 40.73° ± 0.48°; F(1,10) = 7.6, p = 0.02, η²ₚ = 0.43), but without significant Gravity × Direction interactions (F(1,10) = 0.45, p = 0.52). In contrast to the increase in stimulation amplitude in weightlessness, eye movement amplitudes were 32% smaller in microgravity (1 g = 24.36 ± 1.48°; 0 g = 16.56 ± 1.26°) (Figure 3E), resulting in a significant reduction in vertical amplitude gain in weightlessness (1 g: 0.62 ± 0.038; 0 g: 0.40 ± 0.030) (Figure 3D). This gravity effect also interacted with a directional effect on ocular displacement amplitude (Gravity × Direction interactions: F(1,10) = 15.04, p = 0.003, η²ₚ = 0.60) and amplitude gain (Gravity × Direction interactions: F(1,10) = 7.51, p = 0.021, η²ₚ = 0.43). Post-hoc analysis revealed a slight difference (5%) in eye amplitude between forward and backward eye movements in microgravity only (0 g backward: 16.99 ± 1.78°, 0 g forward: 16.13 ± 1.87°, p = 0.0086; 1 g backward: 24.23 ± 2.11°, 1 g forward: 24.49 ± 2.20°, p = 0.37), whereas the gravity effect remained significant and large for both forward and backward rotations (decrease of 30% with weightlessness backward and 34% forward, both with p < 0.001). Similarly, the amplitude gain was larger backward than forward in microgravity (0 g backward: 0.41 ± 0.04, 0 g forward: 0.39 ± 0.04, p = 0.0033; 1 g backward: 0.61 ± 0.05, 1 g forward: 0.63 ± 0.06, p = 0.13) whereas the gravity effect was significant in both direction (decrease of 33% with weightlessness backward and 39% forward, both with p < 0.001).

The Gravity × Direction interaction was also observed for head peak velocity (F(1,10) = 7.16, p = 0.023, η²ₚ = 0.42). Post-hoc analyses revealed that directional effects were small but present in microgravity only, with head sway velocity being 4% faster forward than backward (0 g backward: 67.11 ± 1.32°·s⁻¹, 0 g forward: 69.71 ± 1.36°·s⁻¹, p = 0.0021; 1 g backward: 70.59 ± 1.12°·s⁻¹, 1 g forward: 71.00 ± 1.18°·s⁻¹, p = 0.29). Post-hoc analyses also revealed a slight (5%) but significant effect of gravity on stimulation velocity for backward sway only (p = 0.016), but not for forward sway (p = 0.21), with slightly larger peak velocity in normogravity. In contrast, eye velocity was not affected by direction (Direction: F(1,10) = 0.42, p = 0.53; Gravity × Direction: F(1,10) = 0.03, p = 0.87), but was strongly affected by gravity (F(1,10) = 52.81, p < 0.001, η²ₚ = 0.84). A 22% decrease in eye velocity peak was measured in weightlessness (1 g: 58.29 ± 2.44°·s⁻¹; 0 g: 45.42 ± 2.37°·s⁻¹) (Figure 3F), leading to a significant 42% reduction in vertical VOR velocity gain in weightlessness (1 g: 0.59 ± 0.038; 0 g: 0.34 ± 0.039; F(1,10) = 43.49, p < 0.001, η²ₚ = 0.81) (Figure 3D). VOR velocity gain displayed a significant Gravity x Direction interaction (F(1,10) = 5.72, p = 0.038, η²ₚ = 0.36), however post-hoc analysis were not significant (0g backward: 0.36 ± 0.053, 0g forward: 0.33 ± 0.059, p = 0.093; 1g backward: 0.59 ± 0.054, 1g forward: 0.58 ± 0.054, p = 0.858).

Finally, for the vertical VOR, the eye position phase lead relative to head motion was not affected by gravity (1 g: -1 ± 1.1 ms; 0 g: -15.4 ± 8.3 ms; F(1,10) = 0.005, p = 0.94) nor by direction (Direction: F(1,10) = 1.03, p = 0.33; Gravity × Direction: F(1,10) = 1.96, p = 0.19). Additionally, the median vertical eye position remained stable regardless of gravity conditions (1 g: 2.93° ± 0.36°; 0 g: 2.45° ± 0.41°; F(1,10) = 1.69, p = 0.22).

### Longitudinal Effects on the VOR

For both horizontal and vertical VOR tests, head kinematics (amplitude, velocity, and duration) did not differ between Pre and Post measurements, either on ground or in flight (all p > 0.05). These results confirm that passive rotations were reproducible and comparable across testing sessions.

Horizontal VOR amplitude gain measures showed significant longitudinal effects in flight, with lower values post-parabolas compared to pre-parabolas (Figure 4A). This longitudinal effects was not observed on longer period (pre-flight vs. post-flight). This was revealed by a Ground-Flight x Pre-Post interactions (F(1,10) = 6.39, p = 0.032, η²ₚ = 0.42). Post-hoc analysis revealed that amplitude gain remain stable just after take-off (Ground-Pre = 0.66 ± 0.04, Flight-Pre = 0.66 ± 0.04, p = 0.20), decreased in flight immediately after the parabola (Flight-Pre = 0.66 ± 0.04, Flight-Post = 0.56 ± 0.05, p = 0.001), and was restored to its baseline value after landing (Ground-Pre = 0.66 ± 0.04, Ground-Post = 0.62 ± 0.04, p = 0.89). Horizontal VOR velocity gain showed similar effect with a decrease immediately after the parabolas session (Ground-Flight x Pre-Post interactions : F(1,9) = 6.87, p = 0.028, η²ₚ = 0.43). This gain remained stable just after take-off (Ground-Pre = 0.63 ± 0.04, Flight-Pre = 0.64 ± 0.05, p = 0.21), decreased in flight immediately after the parabola (Flight-Pre = 0.64 ± 0.05, Flight-Post = 0.54 ± 0.06, p = 0.002), and was restored to its baseline value after landing (Ground-Pre = 0.63 ± 0.04, Ground-Post = 0.60 ± 0.05, p = 0.86) . In contrast, VOR latency did not differ between Pre and Post conditions, either on ground or in flight (all p > 0.05).

**Figure 4.**
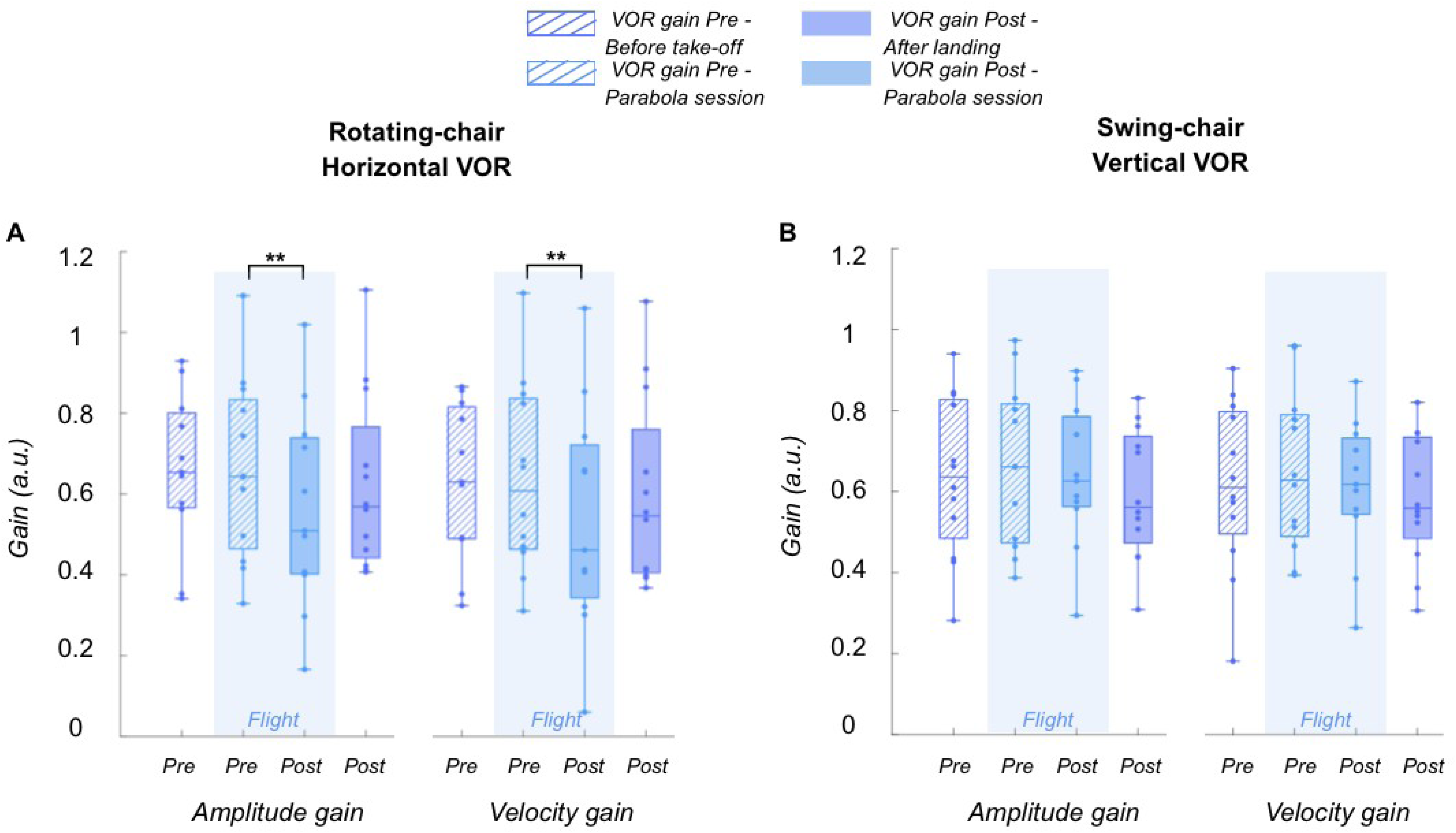
Horizontal (A) and vertical (B) VOR gains depending on time points. Pre- and post-flight measurements (before take-off and after landing) are shown in purple, whereas pre- and post-parabolic session measurements (before parabola 0 and after parabola 30) are shown in blue. Pre-session measurements are represented by dashed boxes, while post-session measurements are represented by filled boxes. Repeated-measures ANOVA with Tukey post-hoc. *, p<0.05, ** p<0.001.

**Figure 5.**
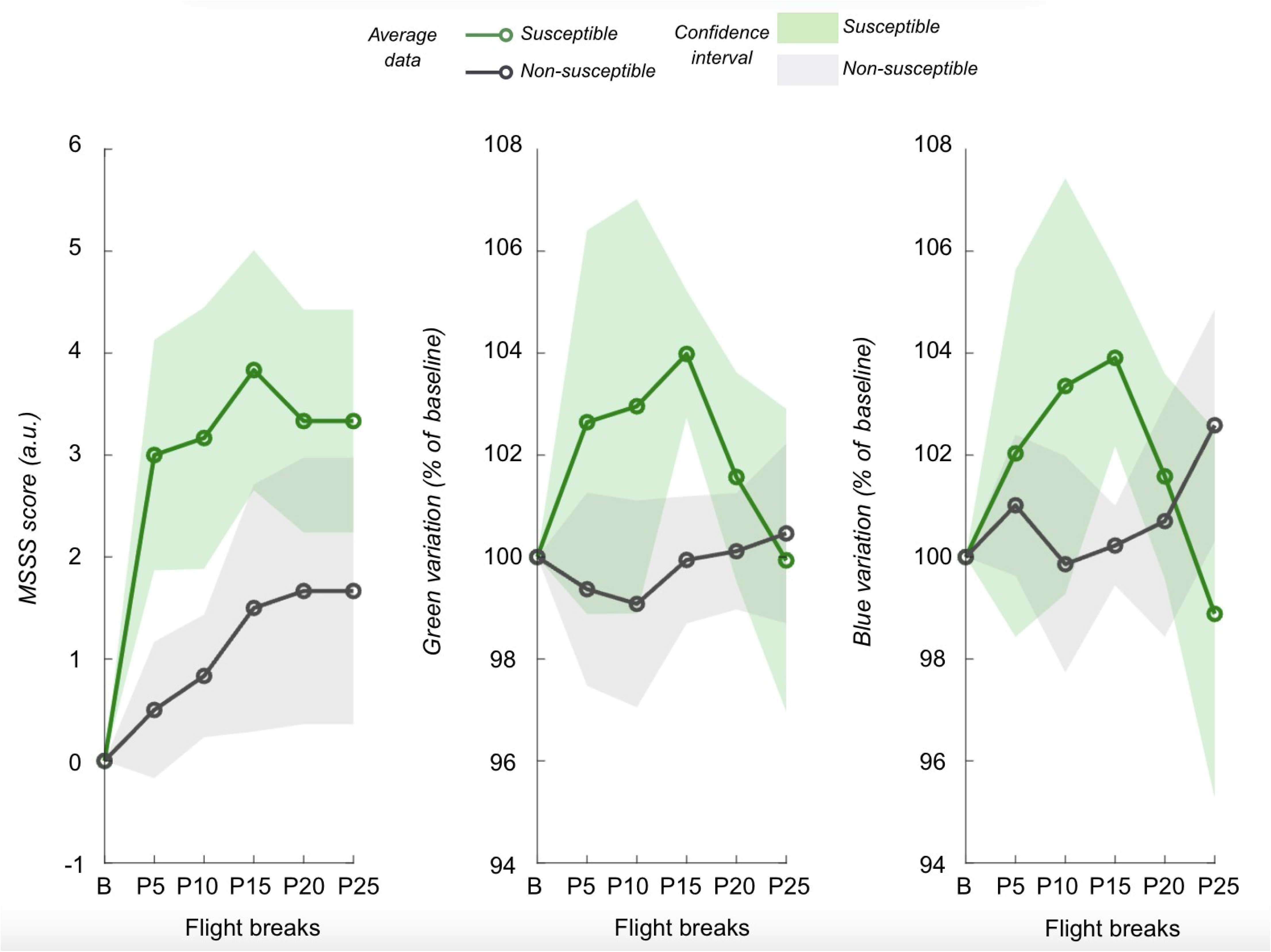
Evolution of subjective sickness ratings (MSSS reports, left panel), facial greenness (Facial green skin color, middle panel) and facial blueness (Facial blue skin color, right panel) across flight breaks. Green curves represent mean values of susceptible (sick) participants, whereas grey curves represent mean values of non-susceptible participants. Confidence intervals (95%) are displayed as translucent areas behind each curve. Six time points are shown, corresponding to the baseline measure (B) and the five in-flight breaks, occurring every five parabolas (P + number of parabola completed). Facial skin color was measured during each break, whereas MSSS scores were recorded at the end of each parabola. MSSS scores were averaged across the two parabolas immediately preceding and the two parabolas immediately following each break. For the Green and Blue scores, percentage changes relative to baseline (100%) values are presented.

Vertical VOR amplitude gain remained unchanged across Pre- and Post-flight and ground measurements (Ground-Pre = 0.64 ± 0.04, Flight-Pre = 0.66 ± 0.04, Flight-Post = 0.64 ± 0.04, Ground-Post = 0.59 ± 0.03; Pre-Post : F(1,10) = 1.08, p = 0.32; Ground-Flight : F(1,10) = 0.77, p = 0.40; Ground-Flight x Pre-Post : F(1,10) = 0.06, p = 0.82) (Figure 4B). Similarly, velocity gain were robust to Pre-Post and Ground-flight effects (Ground-Pre = 0.61 ± 0.04, Flight-Pre = 0.65 ± 0.04, Flight-Post = 0.61 ± 0.03, Ground-Post = 0.58 ± 0.03; Pre-Post : F(1,10) = 1.32, p = 0.28; Ground-Flight : F(1,10) = 0.53, p = 0.48; Ground-Flight x Pre-Post : F(1,10) = 0.03, p = 0.86). Vertical VOR latency remain also unaffected by Pre-Post and Ground-flight effects (all p > 0.05).

### Motion sickness evolution and relationship with VOR performance

We aimed to investigate potential relationships between canal sensitivity, vestibular ponderation, and motion sickness, assessed both subjectively (questionnaires and rating scales) and objectively (skin colorimetry changes). Baseline VOR gains corresponded to the VOR gains measured on the ground prior to takeoff.

Participants reporting greater post-flight symptoms exhibited larger changes in both the green and blue facial color components. Specifically, changes in total SSQ scores were positively correlated with changes in the green component between pre-and post-flight measurements (ρ = 0.736, p = 0.006, n = 12) and with changes in the blue component (ρ = 0.840, p = 0.0006, n = 12). This correlation was also observed when comparing changes in skin color with early in-flight reports of motion sickness symptoms using the MSSS scale. Specifically, MSSS scores obtained at the first break were significantly correlated with changes in the green score (ρ = 0.804, p = 0.002, n = 12) and the blue score (ρ = 0.651, p = 0.022, n = 12). Skin color modifications predominantly occurred during this early phase of the flight, with variations in skin color between baseline and mid-flight (third break) being significantly associated with early motion sickness severity. No significant correlations were found for the red component (all p > 0.05).

Regarding vestibular ponderation, positive correlations were observed between changes in horizontal VOR amplitude gain from pre- to post-flight and the evolution of motion sickness symptoms throughout the flight, with smaller canalar reweighting in sicker participants (SSQ total score difference: ρ = 0.661, p = 0.027, n = 11; MSAQ total score difference: ρ = 0.727, p = 0.015, n = 11) . However, those correlations were not significant for horizontal velocity gain (SSQ total score difference: ρ = 0.410, p = 0.214, n = 11; MSAQ total score difference: ρ = 0.500, p = 0.121, n = 11). Moreover, with respect to canal sensitivity, participants with lower baseline horizontal VOR gains experienced more severe symptoms early in the flight, accompanied by greater increases in facial greenness and blueness (Table 1). For example, higher symptom severity, as measured by MSSS scores at the first break, was associated with lower baseline horizontal gains (amplitude gain: ρ = −0.734, p = 0.010, n = 11; velocity gain: ρ = −0.633, p = 0.037, n = 11). Similar relationships were observed at the second break. Likewise, baseline horizontal VOR gains were negatively correlated with facial color components measured at the first break. Participants with lower baseline horizontal gains exhibited greater facial greenness and blueness (Table 1). Similar correlations were observed at the second break. These relationships between baseline horizontal VOR gain and symptom severity or facial color changes were not observed for the subsequent MSSS assessments. In addition, the red facial color component was not correlated with baseline canal sensitivity. Finally, baseline vertical VOR gains were not associated with symptom severity or with changes in facial greenness or blueness.

**Table 1.** Correlation matrix between VOR gains, facial skin color and subjective sickness. Spearman correlations. Significant correlations are displayed in bold characters (p<0.05)

| Variables | Horizontal VOR |  | Vertical VOR |  |
| --- | --- | --- | --- | --- |
|  | Baseline Amplitude Gain | Baseline Velocity Gain | Baseline Amplitude Gain | Baseline Velocity Gain |
| MSSQ total score | **$\rho = -0.645$, $p = 0.018$, $n = 11$** | $\rho = -0.564$, $p = 0.071$, $n = 11$ | $\rho = -0.153$, $p = 0.634$, $n = 11$ | $\rho = -0.296$, $p = 0.350$, $n = 11$ |
| SSQ total score difference | $\rho = -0.401$, $p = 0.222$, $n = 11$ | $\rho = -0.324$, $p = 0.332$, $n = 11$ | $\rho = 0.049$, $p = 0.879$, $n = 11$ | $\rho = -0.078$, $p = 0.811$, $n = 11$ |
| MSSS break 1 score | **$\rho = -0.734$, $p = 0.010$, $n = 11$** | **$\rho = -0.633$, $p = 0.037$, $n = 11$** | $\rho = -0.021$, $p = 0.948$, $n = 11$ | $\rho = -0.162$, $p = 0.615$, $n = 11$ |
| MSSS break 2 score | **$\rho = -0.712$, $p = 0.014$, $n = 11$** | $\rho = -0.598$, $p = 0.052$, $n = 11$ | $\rho = -0.091$, $p = 0.778$, $n = 11$ | $\rho = -0.200$, $p = 0.533$, $n = 11$ |
| Green break 1 score | **$\rho = -0.747$, $p = 0.008$, $n = 11$** | **$\rho = -0.733$, $p = 0.010$, $n = 11$** | $\rho = -0.319$, $p = 0.313$, $n = 11$ | $\rho = -0.322$, $p = 0.307$, $n = 11$ |
| Blue break 1 score | **$\rho = -0.662$, $p = 0.027$, $n = 11$** | **$\rho = -0.665$, $p = 0.029$, $n = 11$** | $\rho = -0.133$, $p = 0.680$, $n = 11$ | $\rho = -0.133$, $p = 0.680$, $n = 11$ |
| Green break 2 score | **$\rho = -0.688$, $p = 0.019$, $n = 11$** | **$\rho = -0.647$, $p = 0.031$, $n = 11$** | $\rho = -0.410$, $p = 0.186$, $n = 11$ | $\rho = -0.438$, $p = 0.155$, $n = 11$ |
| Blue break 2 score | **$\rho = -0.764$, $p = 0.009$, $n = 11$** | **$\rho = -0.700$, $p = 0.021$, $n = 11$** | $\rho = -0.217$, $p = 0.499$, $n = 11$ | $\rho = -0.245$, $p = 0.444$, $n = 11$ |

## DISCUSSION

We investigated the gravito-inertial effects on vertical and horizontal canal reflexes, as well as their potential evolution in parallel with the emergence of space motion sickness. Based on previous literature, we hypothesized that the vertical VOR would be affected due to its alignment with the gravitational axis. Our results demonstrated that both vertical and horizontal VOR performance decreased in microgravity; however, only the horizontal VOR exhibited a post-flight decrease, which rapidly reversed in normogravity.

The most striking result concerns the effect of weightlessness on the VOR, with a reduced gain observed under this condition compared to the Earth gravitoinertial environment. This supports the idea that studies observing unaffected horizontal reflexes after a few days in microgravity—during or after orbital flight—(Berthoz et al., 1986; Benson & Vieville, 1986; Thornton et al., 1989; Grigorova & Kornilova, 1996; Clarke et al., 2000) reflect an adaptive response to weightlessness rather than a canalar resilience to this context. Our results showed that effects were similar with horizontal and vertical stimulations. Vertical VOR has been investigated far less frequently in previous studies, largely because of the technical constraints, including the need to align the rotation axis with the ears, generate sufficient forces to move the apparatus and reproduce oscillatory motion across gravitoinertial conditions. Thus, the available datas on vertical VOR performance in space originated from studies using active head movements in astronauts (Dai et al., 1994, 1998), which did not showed gravitational effect after a few days in space.

Our results revealed that the significant gravitational effect observed was confounded by an interaction with the direction of pitch oscillation. Specifically, gain was higher during backward motion than during forward motion in weightless only. However, backward stimulations were also characterized by larger amplitude and lower velocity, likely reflecting our inability to reproduce perfectly symmetrical displacements in microgravity due to technical constraints. Although the effect of direction reached statistical significance, its magnitude was very small, with only a 0.7% difference between backward and forward motions in weightless. In contrast, the influence of gravity on ocular amplitude was substantially larger than any directional effect, with a 32% decrease in microgravity, regardless of direction. Consequently, the reduction in gain observed in microgravity cannot be attributed primarily to negligible differences in stimulation kinematics.

Two main hypotheses may be proposed to explain the observed reduction in VOR performance. The first mechanism involves alterations in semicircular canal fluid kinematics. Several biomechanical studies of semicircular canal function suggest that these structures are sensitive to pressure (Muller, 2017; Yamauchiet al., 2002). Changes in inner-ear pressure could generate compressive force gradients within the semicircular canal endolymph, thereby perturbing the dynamics of inertial fluid motion (Rabbitt, 2019). Such alterations in endolymph dynamics would affect cupular deflection and, consequently, impair canal signal transduction in weightless conditions. One potential driver of these changes is the cephalad fluid shift that occurs during the initial phase of exposure to microgravity, which is thought to increase intracranial pressure (Avan et al., 2018). Consequently, semicircular canal mechanics during the 0 g phases of parabolic flight may be altered by these pressure changes, potentially contributing to the reduction in VOR performance observed in microgravity.

The second hypothesis involves the central integration and processing of vestibular signals by internal models within the brainstem and cerebellum. Although the VOR primarily relies on a short three-neuron reflex arc, it is well established that additional, more complex neural pathways can modulate this response (Green & Angelaki, 2010), including projections with rostral fastigial nucleus and the flocculonodular lobe (Brooks & Cullen, 2009). Several internal model theories propose that the central nervous system relies on gravito-inertial information to estimate body orientation and self-motion as well as to generate appropriate motor commands (Angelaki & Cullen, 2008; Green & Angelaki, 2010; Cullen, 2023). Consequently, the novelty of weightlessness may impair the functioning of these internal models, which must adapt and recalibrate motor commands to the new biomechanical constraints of the system. As a result, the oculomotor commands may be inaccurate, leading to under-compensation of head motion and, ultimately, a reduction in VOR gain. Furthermore, in this experiment, the lack of visual feedback prevented error identification and recalibration of the internal models.

Our second major finding was a post-flight reduction in horizontal VOR gain, which returned to baseline shortly after landing. To our knowledge, no previous studies have directly compared VOR gain before and after parabolic flight—a comparison typically conducted after orbital flight. A reduction of active VOR gain has been depicted post-mission (Clarke et al., 2000 ; Clément et al., 2019), also found using post-rotatory nystagmus (Oman, 1996). In contrast, earlier work by Thornton et al. (1989) did not reveal any significant changes in horizontal VOR gain post-mission. Taken together, a majority of studies suggest a decrease in VOR gain following adaptation to microgravity, consistent with our observations after parabolic flight. Importantly, these studies also indicate that such reductions are transient and recover rapidly, as appears to be the case in our protocol. Nevertheless, we suggest that the observed VOR gain decrease in our protocol reflects transient sensory reweighting—with reduced semicircular canal reliance—rather than true adaptation to weightlessness. To compensate for the underdetection of the channel, the VOR gain should actually have been increased after the flight, rather than decreased as observed here. On the other hand, sensory inputs may be selectively reweighted following exposure to sensory conflicts. Based on Bayesian theory of multisensory integration, sensory signals that are deemed less reliable relative to others are downweighted in order to reduce overall perceptual uncertainty (Knill & Pouget, 2004). In a previous study, we demonstrated that exposure to parabolic flight decreased the gain of the visual optokinetic nystagmus, while leaving the performance of otolith-driven reflexes unaffected (Bonnard et al., 2025). In this previous experiment, we also observed a strong trend toward a pre–post effect in horizontal VOR suppression, with reduced VOR gain (i.e., more effective suppression) following flight. This trend, together with our new results, suggests that semicircular canal inputs may be selectively downweighted after exposure to parabolic flight.

One possible explanation for these down-weightings lies in the sensory context experienced during parabolic flight. Under microgravity conditions, both otolithic and proprioceptive signals consistently indicate free fall. In contrast, neither vertical optic flow nor adequate head movements are perceived. Consequently, visual and canalar signals become incongruent with otolithic and proprioceptive information. From an evolutionary perspective, otolithic cues provide robust and stable information about gravito-inertial forces, whereas inputs from the semicircular canals and visual system are more variable and context dependent. As a result, visual and canalar signals may be considered less reliable under these conditions and therefore assigned lower weights during multisensory integration than otolithic inputs.

In addition, in all our conditions, we observed negative VOR latencies, suggesting that eye movements preceded head movements. One possible explanation is the presence of anticipatory eye movements driven by prediction of the upcoming head motion. Given that head oscillations were performed with fixed amplitudes and relatively constant frequencies, participants may have been able to internalize the movement rhythm and anticipate direction changes. However, the relatively large standard errors associated with these latency estimates indicate substantial inter-individual variability, suggesting that such anticipatory behavior was not consistently present across participants. These findings should also be interpreted with caution in light of the methodology used to estimate latency. The current approach relies on aligning the entire eye and head movement signals, whereas the temporal delay of interest is primarily determined by the onset of each movement. Consequently, the observed negative latencies may partly reflect methodological limitations rather than genuine anticipatory responses.

Finally, in terms of objective assessment of motion sickness severity, we replicated our previous findings regarding facial greenness during parabolic flight (Bonnard et al., 2026c). Participants who reported more severe symptoms exhibited the largest increases in the green and blue components of facial skin color, especially during early flight. The absence of a corresponding increase in the red component, together with the predominance of the green and blue changes, supports the hypothesis of facial vasoconstriction. Such vasoconstriction may enhance the relative contribution of skin pigments and deoxygenated blood to the overall appearance of the skin, thereby producing the characteristic facial discoloration associated with motion sickness.

Furthermore, we identified significant correlations between baseline horizontal semicircular canal sensitivity, subjective symptoms, and facial color changes during the early phase of flight. Participants with lower baseline horizontal VOR gains experienced more severe motion sickness and exhibited greater facial greenness and blueness during the first half of the flight. In contrast, no such relationships were observed for motion sickness severity at later time points or for post-flight symptom evolution, demonstrating the motion sickness fate is mostly established during the first 10^th^ parabola. General motion sickness susceptibility also appeared to be associated with baseline horizontal VOR gain. Interestingly, these associations were specific to horizontal VOR gain, whereas vertical VOR gain showed no correlation with either subjective or objective measures of motion sickness. This finding is somewhat unexpected, given that pitch head movements are typically considered more provocative than yaw movements and are more likely to generate otolitho-canal conflicts. Our results, however, did not support the idea of a predominant contribution of vertical canal function to in-flight susceptibility. Earlier studies have mainly focused on vestibular dynamics and velocity storage mechanisms, whereas the present study examined baseline VOR gain. Nevertheless, the interpretation remains consistent, as both approaches suggest that reduced semicircular canal sensitivity or impaired vestibular performance may contribute to increased susceptibility to motion sickness. Clément and Reschke (2018) found no relationship between baseline horizontal VOR gain and motion sickness severity induced by parabolic flight, although they reported that participants experiencing greater symptoms exhibited longer VOR time constants. Furthermore, other studies have shown that larger ocular deviations relative to the gravito-inertial vector during vestibular stimulation are associated with increased motion sickness susceptibility (Dai et al., 2003; Cohen et al., 2003). These findings suggest that vestibular dynamics and velocity storage mechanisms, rather than VOR gain alone, could play a more prominent role in determining individual susceptibility to motion sickness.

To sum up, semicircular canal performance is impaired in microgravity. Although only horizontal VOR performance was found to decrease following flight exposure— likely due to a temporary discrediting of canal cue reliability—this impairment appears to be rapidly reversible. Furthermore, canal sensitivity seemed associated with motion sickness susceptibility during flight. Finally, our findings confirm facial greenness as a relevant objective marker of sickness severity during parabolic flight and potentially of space-like motion sickness.

## FUNDINGS

The authors acknowledge financial support from the Centre national d’études spatiales (CNES), France (ROR: https://ror.org/04h1h0y33), within the framework of the GRAVISMS project. This study also received financial support from the French government in the framework of the University of Bordeaux’s IdEx “Investments for the Future” program / GPR BRAIN_2030.

## DISCLOSURES

No conflicts of interest, financial or otherwise, are declared by the authors.

## Notes

### Competing Interest Statement

The authors have declared no competing interest.

